# High-throughput confocal airy beam oblique light-sheet tomography of brain-wide imaging at single-cell resolution

**DOI:** 10.1101/2023.06.04.543586

**Authors:** Xiaoli Qi, Rodrigo Muñoz-Castañeda, Arun Narasimhan, Liya Ding, Xin Chen, Corey Elowsky, Jason Palmer, Rhonda Drewes, Jianjun Sun, Judith Mizrachi, Hanchuan Peng, Zhuhao Wu, Pavel Osten

**Affiliations:** Cold Spring Harbor Laboratory, Cold Spring Harbor, NY, 11724, USA; National Institute of Biological Sciences, Beijing (NIBS), Beijing, 102206, China; Institute for Brain and Intelligence, Southeast University, Nanjing, 210096, China; Appel Alzheimer’s Disease Research Institute, Feil Family Brain and Mind Research Institute, Weill Cornell Medicine, New York, NY, 10021, USA

## Abstract

Brain research is an area of research characterized by its cutting-edge nature, with brain mapping constituting a crucial aspect of this field. As sequencing tools have played a crucial role in gene sequencing, brain mapping largely depends on automated, high-throughput and high-resolution imaging techniques. Over the years, the demand for high-throughput imaging has scaled exponentially with the rapid development of microscopic brain mapping. In this paper, we introduce the novel concept of confocal Airy beam into oblique light-sheet tomography named CAB-OLST. We demonstrate that this technique enables the high throughput of brain-wide imaging of long-distance axon projection for the entire mouse brain at a resolution of 0.26 μm × 0.26 μm × 1.06 μm in 58 hours. This technique represents an innovative contribution to the field of brain research by setting a new standard for high-throughput imaging techniques.

## Introduction

As brain science continues to advance, countries around the world are increasingly investing resources into cutting-edge research in the field. Among the key research directions in brain science is the drawing of microscopic brain maps, which serve as a fundamental for achieving a comprehensive understanding of the brain. One of the American Brain Project’s primary ventures, BRAIN Initiative Cell Census Network (BICCN), is targeted to mapping of motor cortex cells. Just as gene sequencing is highly dependent on sequencing tools, the drawing of microscopic brain maps is highly dependent on large-volume, high-resolution, and automated imaging tools. Previously, micro wide mapping^1^. Following MOST’s success, serial two-photon tomography (STPT) was developed^2^. STPT enabled a range of research on mapping long-range connectivity of thalamic projections, cortex-striatum connections, and region-to-region connectivity through the Allen Institute for Brain Science Mouse Connectivity Project. This project has successfully assayed the brain-wide projections of genetically identified cell populations in hundreds of distinct brain regions, as previously -optical sectioning tomography (MOST) was successfully utilized to fully automated brain-reported^3^. Subsequently, the drawing of microscopic brain maps developed rapidly, and at the same time, the demand for high-throughput increased exponentially.

Light-sheet fluorescence microscopy(LSFM), together with the tissue clearing technique, significantly extends the penetration depth of light and enables imaging of large volumes with faster speed and less photonbleaching^4–6^. Those features made light-sheet microscopy potential useful in neuroanatomy^7^. However, most light-sheet microscopy is limited by low resolution (cellular), undersized imaging volume, and poor compatibility of different clearing techniques with different refractive index (RI)^8^. To overcome these limitations, this paper first proposes the concept of confocal Airy beam imaging to light-sheet imaging and develops confocal Airy beam oblique light-sheet tomography (CAB-OLST). CAB-OLST provides excellent image quality over large fields-of-view (FOV), a large imaging volume, and a simple and wide RI at a wide range of 1.33-1.55. And it enables fully automated high resolution with high-throughput imaging of fluorescently labeled mouse brains. The long-distance projection data of the entire mouse brain was successfully obtained using CAB-OLST, making it the world’s fastest technique with single-cell resolution (0.26 μm × 0.26 μm × 1.06 μm).

## Results

### High-throughput, high SNR light-sheet tomography

In the field of high-resolution imaging techniques, CAB-OLST has emerged as an efficient, automated solution for high-throughput imaging with high signal-to-noise ratio (SNR) (**Tab. 1**). CAB-OLST works as below (**Fig. 1a**). An agar-embedded cleared mouse brain is mounted in an oil bath on integrated x-y-z stages with an inverted selective plane illumination microscopy (SPIM) structure, and imaging parameters are then set, following which the imaging system operates automatically for a duration ranging from a few hours to a few days, depending on the sample size and the sampling parameters (**Tab. 2**).

**Table 1.** Throughput of whole mouse brain imaging

| | voxel resolution<br>($\mu\text{m}$) | total imaging<br>(hours) |
| --- | --- | --- |
| Our | $0.26 \times 0.26 \times 1.06$ | 29/58 |
| Mouse light<sup>18</sup> | $0.3 \times 0.3 \times 1.0$ | ~168 |

**Figure 1.**
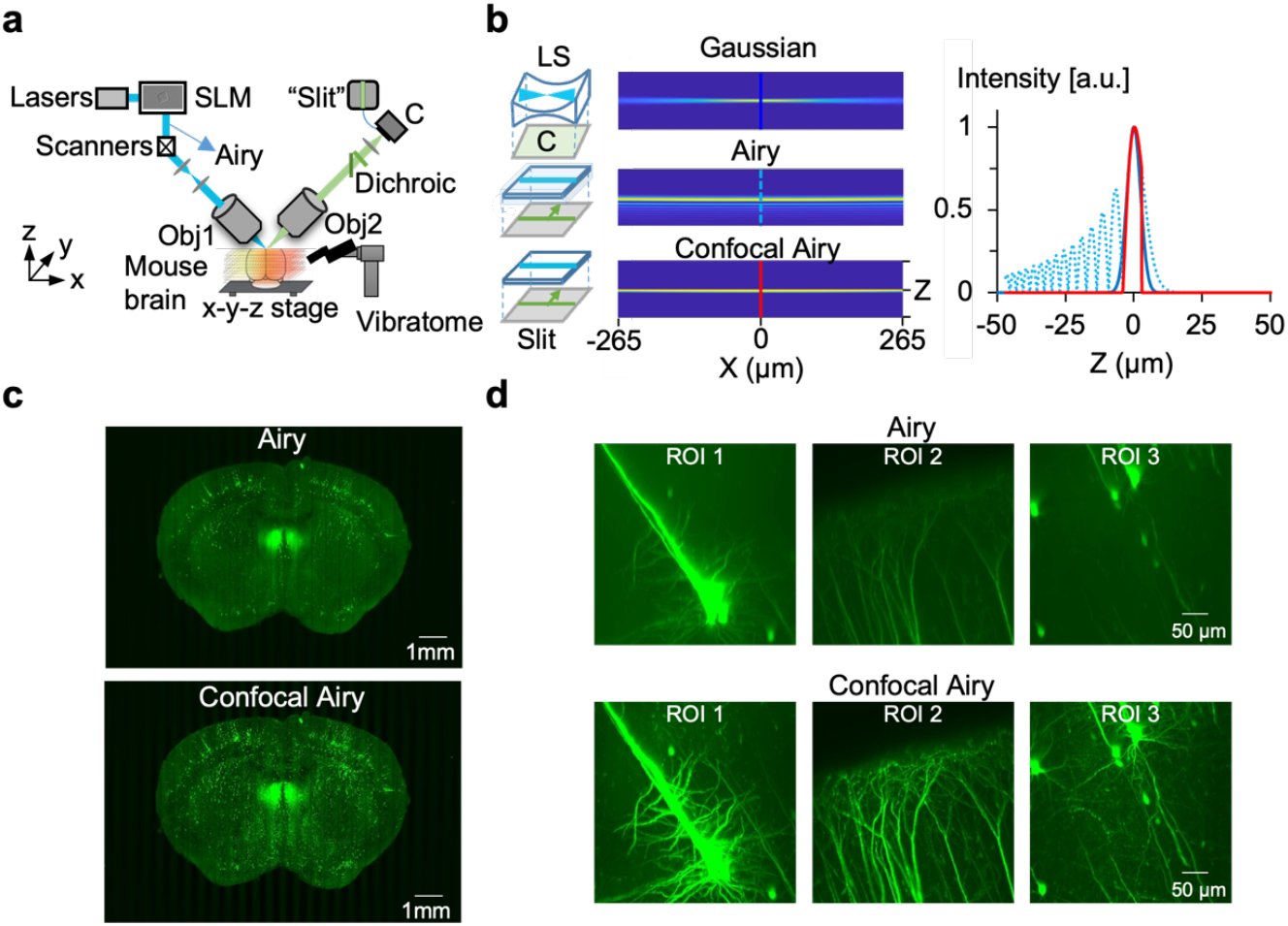
CAB-OLST. a) System principal diagram. Spatial light modular shapes the Gaussian beam to Airy beam to extend the depth of field (DOF) of a SPIM light-sheet (LS) structure. An agar-embedded cleared mouse brain is imaged with the movement of the x-y-z stage, then moves to a vibratome location to slice the coronal plane. b) Light-sheet formation from standard Gaussian beam-based LS, airy beam-based LS and straight Airy beam-based LS. A straight Airy beam extends and uniforms the DOF compared to the Gaussian beam, and side lobes are removed by a confocal slit. c, d) Image comparison with Slit Confocal Airy LS and standard Airy LS. Applying Slit Confocal to the straight airy beam improves optical sectioning ability without deconvolution.

**Table 2.** Throughput of whole brain imaging of CAB-OLST

| | camera<br>height<br>($\mu\text{m}$) | camera<br>width<br>($\mu\text{m}$) | voxel resolution<br>($\mu\text{m}$) | total<br>imaging<br>(hours) |
| --- | --- | --- | --- | --- |
| Airy LS |  |  |  |  |
| Obj.4 pair | 756 | 756 | $0.37 \times 0.37 \times 1.77$ | 10 |
| Obj.7 pair | 530 | 530 | $0.26 \times 0.26 \times 1.77$ | 21 |
| Obj.7 pair | 530 | 530 | $0.26 \times 0.26 \times 1.06$ | 29 |
| Obj.7 pair | 530 | 530 | $0.26 \times 0.26 \times 0.71$ | 52.5 |
| Airy confocal LS |  |  |  |  |
| Obj.4 pair | 756 | 756 | $0.37 \times 0.37 \times 1.77$ | 20 |
| Obj.7 pair | 530 | 530 | $0.26 \times 0.26 \times 1.77$ | 42 |
| Obj.7 pair | 530 | 530 | $0.26 \times 0.26 \times 1.06$ | 58 |
| Obj.7 pair | 530 | 530 | $0.26 \times 0.26 \times 0.71$ | 88.5 |

During imaging, the x-y-z stages move the brain below the objective lens, and an airy beam scanned light-sheet generates oblique optical section that are subsequently imaged as a mosaic of fields of view (FOVs) to create the superficial volume of the whole brain. After, mechanical sectioning is implemented to remove the imaged tissue section using the built-in vibratome. The optical and mechanical sectioning steps continue in repetition until the dataset is collected. To illuminate a larger field-of-view uniformly, an airy beam scanned light-sheet to extend the depth of field (DOF) 6∼10 times^9,10^, comparing to the typical Gaussian beam (**Fig. 1b**). It increases system throughput while maintaining a thin thickness of the light sheet. And a straight airy beam that can be formed by rotating the angle of curved propagation-invariant Airy beam^11^. Furthermore, a virtual slit was added to remove the side lobes of the airy beam without the need for deconvolution.

To demonstrate increased SNR through background suppression, we used Thy1-GFP (line M) mice which express GFP mainly in hippocampal and cortical pyramidal neurons with complete morphology. **Fig. 1c, d** shows the imaging of the same coronal section with slit confocal Airy beam light-sheet and typical Airy beam light-sheet. Applying slit confocal to the straight airy beam improves optical sectioning ability, and there is no need for time-consuming deconvolution. Further, **Fig. 2a, b** demonstrates the quantitative comparison in different signal locations (soma, neurites). The contrast improves for both sites, 4.7 times improvement in neurites while 1.2 times in soma (n=5). It reveals that a confocal airy light-sheet increases SNR in all locations and might be vital for having complete morphology.

**Figure 2.**
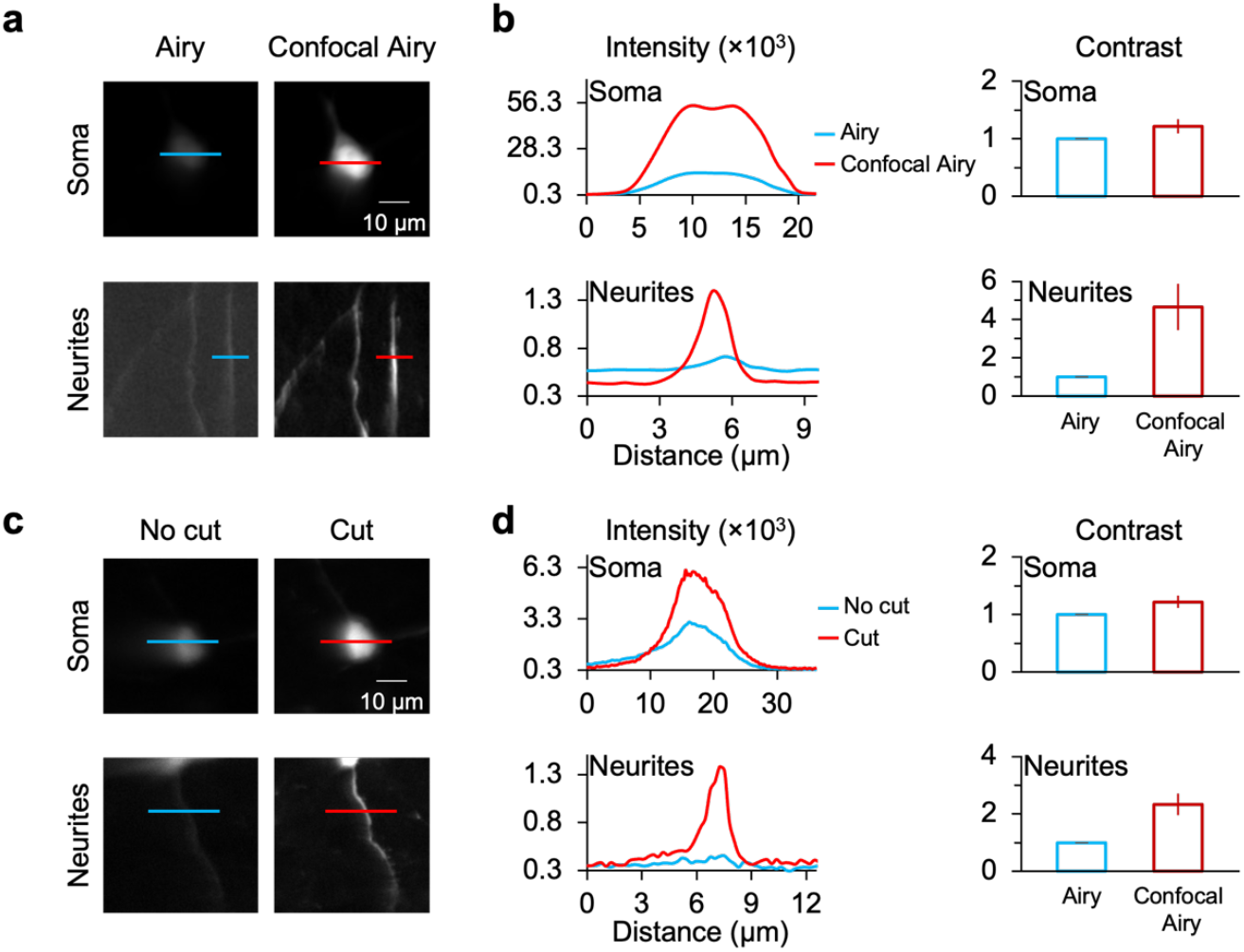
Quantitative comparison shows high SNR of CAB-OLST. a,b) Quantitative comparison with Slit Confocal Airy LS and standard Airy LS in different signal locations(soma, dendrite). c,d) Image comparison with standard LS (no cut) and cut LS. The cut LS can achieve high quality and SNR across the whole brain. While the images of standard depth LS might degrade in the deeper position, making fine structure invisible.

Although an advanced tissue clearing method (U Clear, unpublished, Zhuhao Wu) is applied here, we still can benefit from the strategy of combing optical sectioning and mechanical sectioning. First, our cutting methods can achieve the same high-quality images through all depths of large volumes. The second fine structure with low SNR can still be detectable in the deeper location. **Fig. 2c, d** demonstrates comparisons with common light-sheet (no cut) and our serial sectioning light-sheet (cut). The cut light-sheet can achieve high quality and signal-noise-ratio across the whole brain. While the images of typical depth light-sheet degrade in the deeper position, making fine structure invisible. The fully automated imaging and cutting design has minimal detrimental effects and shows an inherent alignment feature in 3D. Besides, a new pair of multi-immersion objectives (special optics, Inc.) replace the water-based objectives to match a better refractive index at a wide range of 1.33-1.55. Also, benefiting from the long working distance and unique design geometry, this scope has a 2 mm to 5 mm co-working distance which is not typical for standard objective pair. Co-localized multi-channel imaging is also added into the system as well the multi-channels have aligned in 3D of physical system, with no need to align in the post-processing. **Fig. 3** shows the measurement results of lateral and axial point spread function. **Fig. 4** shows Quantitative statistics of PVH projections in brain wide with a single-cell resolution.

**Figure 3.**
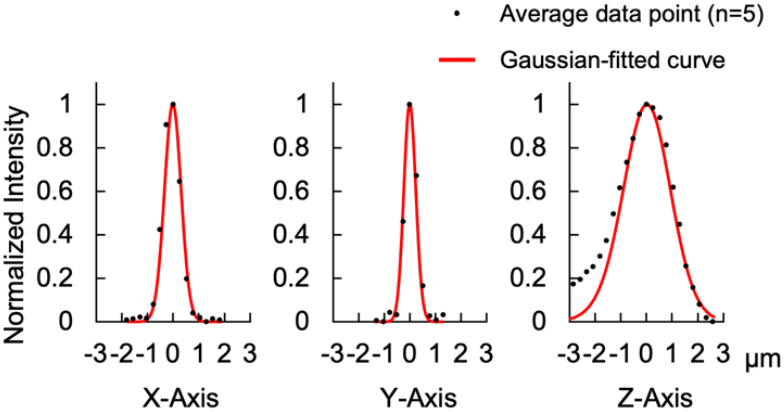
Measurement of lateral and axial point spread function. X: 0.76 μm ± 0.06 μm; Y: 0.53 μm ± 0.04 μm; Z: 2.20 μm ± 0.35 μm.

**Figure 4.**
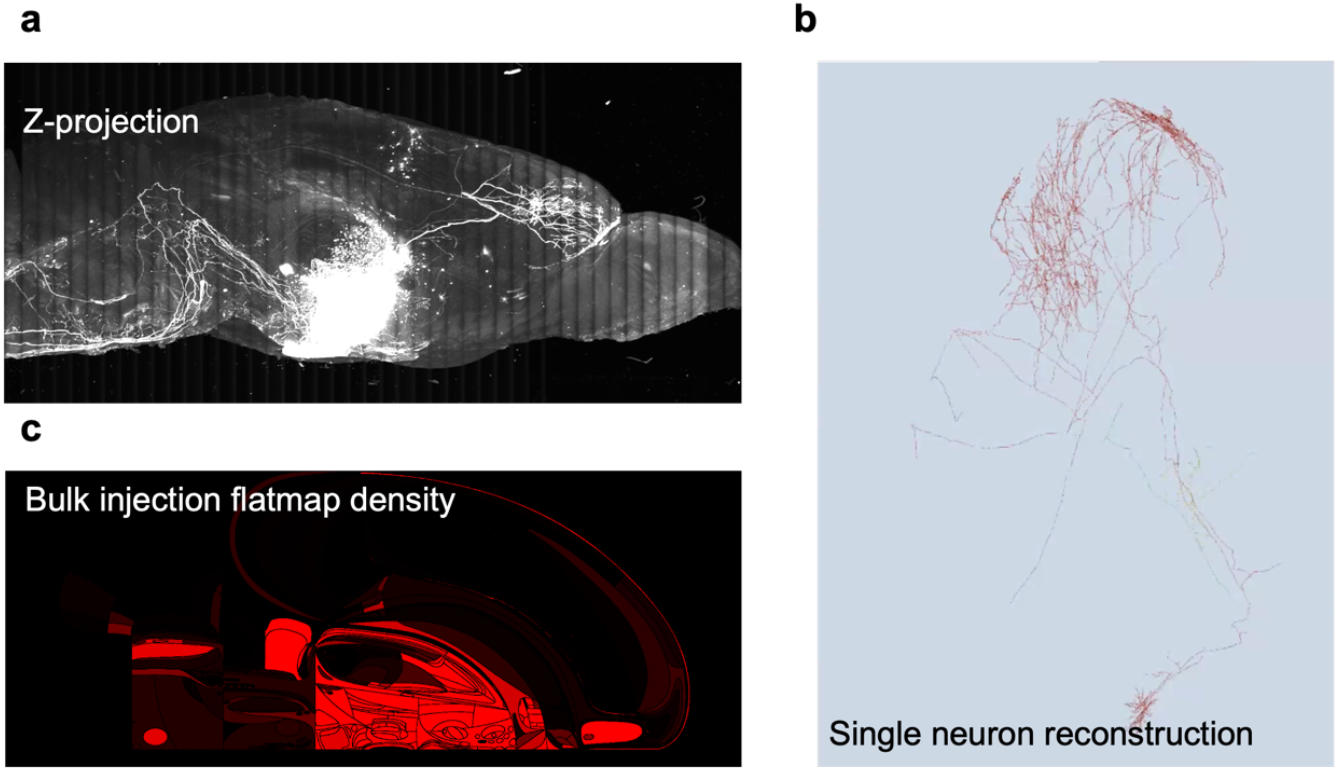
Quantitative statistics of brain-wide PVH projections with a single-cell resolution. a. Maximum Intensity Projection (MIP) via Z axis of AVPtm brain. b. bulk projection flatmp density c. single neuron reconstruction.

## Discussion and Conclusion

In contrast to standard light-sheet microscopy, CAB-OLST has many advantages. Firstly, standard light-sheet microscopy employs a Gaussian beam and has a tradeoff of throughput and resolution in large volume^8^. While this method uses an airy beam scanned light-sheet replacing the Gaussian beam to extend the depth of field (DOF) to generate a thinner and larger FOV to achieve higher throughput. Fast axial scanning light-sheet microscopy^12,13^ achieves large FOV by axially scanning a small Gaussian FOV. Extending the DOF methods like CAB light-sheet effectively transfers the 2D frame rate to the volume rate, no need for extra relay optics and synchronized mechanism, especially suitable for higher throughput for sparsely expressed samples^14^. Secondly, although tissue clearing was applied to highly scatted biological tissue, light attenuation and aberration through penetration depth are still a hurdle to see the fine structure without compromised effective spatial resolution^7^. Instead, by combing optical sectioning and mechanical sectioning, CAB-OLST only images less than 1mm deep into the tissue to eliminate most quality degrades related to penetration depth. It achieved less optical aberrations, higher actual axial resolution, and uniform contrast through all depths of large volumes. Third, benefitting from the structure of inverted SPIM or oblique light sheet^15,16^, the sample size is not limited by the sample chamber or the working distance of the objective lens. It can extend to larger samples like rat brains, monkey brains, or human brains.

Automated block-face imaging integrated tissue sectioning represents a critical role in large-scale neuroanatomy research^2,17^. Compared to STPT^2^ and its advanced version of volumetric two-photon tomography^18^, CAB-OLST is light-sheet-based method which increases imaging speed and signal integration time to boost system throughput and resolution. Compared to OLST^16^, an airy confocal beam scanned light-sheet replaces the Gaussian beam to extend the DOF to generate a uniform and low noise light-sheet. Thus, we have increased axial resolution and SNR with increased system throughput. More, a new pair of multi-immersion objectives replace the water-based objectives to better match cleared samples in a range of 1.33-1.55, plus have 2mm to 5mm co-working distance. Co-localized multi-channel imaging is also added.

In summary, CAB-OLST is the first airy beam-based light-sheet tomography that enables whole mouse brain imaging with high 3D spatial resolution. And it first applies virtual confocal slit into airy beam light-sheet microscopy. To the best of our knowledge, it is also the fastest entire mouse brain imaging system with a resolution 0.26 μm × 0.26 μm × 1.06 μm. Because of the excellent balance of high throughput and high resolution, this method can be a new generation of automated light microscopy tomography to systematically generate brain atlas and facilitate the process of generating microscopic single-neuron projections or mesoscale distribution of mammalian brains.

## Methods

### Mice

Animal procedures were approved by the Cold Spring Harbor Laboratory Animal Care and Use Committee and carried out in accordance with US National Institutes of Health standards. Individual genetic lines include adult (8-10 weeks) wildtype mice of the C57BL/6J (Jackson, stock no: 000664), Oxt-Cre mice (Jackson, stock no: 024234), Avptm-Cre mice (Jackson, stock no: 023530), Thy1-GFP mice (Jackson, stock no: 007788). Mice were maintained on a 12-h light/dark cycle with food and water available ad libitum.

### Viral injection

For anterograde neural labeling, virus AAV.DIO.eGFP. (4.3×10^13^ GC/ml, addgene. stock no. 100043) were diluted 10 times and typically obtained at a tilter of 4.3×10^12^ GC/mL. Mice were anesthetized with 1.5% isoflurane before surgery and then placed in a mouse stereotaxic instrument. The injection was performed using a Nanoject II microinjection device (Drummond Scientific). and delivered the virus to the target areas (AP, - 0.6, - 0.8, - 1.0 mm; ML, + 0.2 mm; DV, + 4.9, + 4.95, + 4. 75mm) of at 150nl min^-1^ correspondingly. All subsequent experiments were performed 6 weeks after virus injection to allow sufficient time for transgene expression.

### Sample preparation

The mice were deeply anesthetized and transcardially perfused intracardially, embedded in melted oxidized agarose, and covalent cross-linking as described^2^. Tissue clearing of was performed using U Clear protocol (unpublished, Zhuhao Wu) for all brains. For Oxt-Cre mice and Avptm-Cre mice, Whole-mount brains, immunostaining was performed.

### Imaging system and software

The system consists of a home-build Airy beam scanned path, an inverted selective plane illumination microscopy (SPIM) (ASI.Inc, SPIM), and a customized vibratome sectioning setup. Laser light from CW Solid State Lasers (OBIS 488nm, 561nm, 640nm, Coherent) is expanded and then fills the aperture of spatial light modular (1920 × 1152, Meadowlark). A Cubic phase modulation introduces a phase delay Δφ“u, v” = 2πα (u^3^ + v^3^), then the illumination beam goes through two 4f relays and scanned galvanometer mirrors (8315K, Cambridge Technology), and reimaging it to the back aperture of the illumination objective (25× NA 0.7, n =1.528, Special optics). The fluorescent signal from the sample is collected by detection objective (either a 25× NA 0.7, n =1.528, Special optics, or 17× NA 0.4, n =1.528). It passes through a dichroic mirror (T565LPXR-UF1, Chroma), an emission filter (ET525/50m and ET655LP) and is directed by a tube lens (f=200mm, Nikon) and onto an sCMOS camera (2048 × 2048 pixels, pixel size 6.5 μm, Andor Zyla 4.2 sCMOS).

Robust mechanical sectioning was achieved by a vibrating blade (Ceramic Blade, Precisionary Instruments) vibratome (Leica, Section Head VT1000S) integrated into the imaging system. The vibration frequency can be set to 0-80Hz and the blade angle to 5-20 degrees, amplitude from 0.2mm to 1.0 mm. In our experiments, we use an amplitude 0.8mm at 60Hz at a blade angle of 15 degrees. To increase reliability, z-planes were overlapping before and after sectioning during a whole-brain dataset.

The instrument was controlled by a manufactory’s provided software named ASI-diSPIM, a plugin in an open-source software Micro-Manager. Combined with ImageJ, Micro-Manager provides full-featured microscope management and image processing package comparable to commercial solutions. A customize shell script was used for handles the timing of stage motion, data acquisition, and vibratome control.

### Instrument operation

Once the brain is positioned under the objectives, and the imaging and sectioning parameters are chosen, the instruments operate in a fully automated mode. The brain is mounted in an oil bath (Cargille Laboratories, RI=1.528, which it equal to U Clear buffer) positioned on the computer-controlled x-y-z stages. After identifying the z position of the brain surface under the objectives, the following parameters are set in the software: FOV size, FOV mosaic size, pixel size, confocal slit width and pixel residence time, laser power, sectioning speed, sectioning frequency, z step for each sectioning cycle and number of z sections. The start of the imaging plane is set below the brain surface to ensure an undisturbed optical sectioning surface. Typically, we imaged 50 μm below the surface (through Z or vertical plane).

The single FOV of 45 degrees of the oblique plane is 530 μm × 530 μm. Motion imaging in the x stage with x step 1∼2.5 μm was implemented to acquire one volume stack in one y stage position. In experiments with the 25× objective, we used the 25 y positions with y step 500 μm to cover one section. Once a section was completed, the same x-y-z stage used for imaging moves the sample from the microscope objective toward a vibratome blade to section the uppermost portion of the tissue (cut thickness = 300 μm). There was 45∼47 sections of mouse brain dataset. The overlaps for Y and Z sections are allowed for post-processing stitching.

### Imaging processing

Full neuronal morphology reconstruction was done by our collaborators at institute for Brain and Intelligence. In brief, trained reconstructors use the Vaa3D suite of tools to complete their reconstructions with semi-auto reconstruction. A final QC-checking procedure is always performed by at least one more experienced annotator using TeraVR who reviews the entire reconstruction of a neuron at high magnification paying special attention to the proximal axonal part or a main axonal trunk of an axon cluster where axonal collaterals often emerge and branches are more frequently missed due to the local image environment being composed of crowded high contrasting structures. To finalize the reconstruction, an auto-refinement step fits the tracing to the center of fluorescent signals. The final reconstruction file (.swc) is a single tree without breaks, loops, or multiple branches from a single point.

## Funding

This work was supported by NIH U01MH114824 and U19MH114821.

## Acknowledgments

We thank Rob Campbell for sharing his design for vibratome.

## Notes

### Competing Interest Statement

The authors have declared no competing interest.

